# Chemoenzymatic Measurement of Cell-surface Glycan in Single-cell Multiomics: LacNAc as an Example

**DOI:** 10.1101/2022.09.01.506123

**Authors:** Wenhao Yu, Xinlu Zhao, Abubakar S. Jalloh, Yingying Zhao, Brandon Dinner, Yang Yang, Shian Ouyang, Yachao Li, Tian Tian, Zihan Zhao, Rong Yang, Mingkuan Chen, Gregoire Lauvau, Zijian Guo, Peng Wu, Jie P. Li

## Abstract

Despite the rich information of a cell’s physiological state encoded in the dynamic changes of cell-surface glycans, methods of capturing glycosylation states at the single-cell level are quite limited. Here we report a chemoenzymatic single-cell N-acetyllactosamine (LacNAc) detection method via tagging the LacNAc with a specific DNA barcode. Compared to the lectin-based glycan detection, the chemoenzymatic labeling does not change the transcriptional status of immune cells and is more compatible with scRNA-seq. Integrated analysis of LacNAc and transcriptome of T cells at a single-cell level reveals that the quantity of cell-surface LacNAc is significantly upregulated in activated CD8^+^ T cells but maintained at the basal level in quiescent CD8^+^ T cells (i.e., naive and central memory T cells). Further analysis confirms that the LacNAc level is positively correlated to the glycolytic activity of CD8^+^ T cells at all statues. Taken together, our study demonstrates the feasibility of chemoenzymatic detection of cell-surface glycan in single-cell RNA sequencing-based multiomics with information of TCR sequence and cell-surface epitopes (i.e., scTCR and CITE-seq) and offers a new way to characterize the biological role of glycan in diversified physiological states.

---

A central objective of immunology is to study the incredible cellular diversity and the developmental trajectories of the immune system, which requires resolving cell identities at high resolution^1^. The revolution of single-cell technologies is paving the way towards this goal, unraveling immune system one cell at a time^2–4^. Among these new technologies, the high-throughput single-cell RNA sequencing (scRNA-seq) has proven invaluable for describing heterogeneous immune cell populations^5–9^. However, analysis of transcripts alone only provides a partial picture of cellular states and functions, which can neither resolve the cell identity comprehensively nor allow the analysis of the connections between transcription and other modalities^10^. From a chemist’s point of view, a cell’s identity would be defined as the entire collection of molecular modalities, including DNAs, RNAs, proteins, lipids and posttranslational modifications, as well as their interconnections within a cell^11^. Recent studies have demonstrated the possibility of simultaneous measurement of multiple molecular patterns in single cells, affording a more complete understanding of the cellular states^12–19^. For example, one of the most widely used single-cell multiomic technique is the cellular indexing of transcriptomes and epitopes by sequencing (CITE-seq)^14^, which layers the cell-surface protein characterizations on top of scRNA-seq data and enables the integrated analysis to adjust the differences between transcript and protein quantifications (**Scheme 1**).

Despite being an important posttranslational modification of membrane proteins, glycosylation has rarely been characterized at the level of single cells. Nevertheless, almost all cell-surface receptors involved in the innate and adaptive immune systems are glycosylated^20^. Certain glycan epitopes have been used to describe particular states of immune cells^21, 22^. For example, sialyl Lewis x (CD15s) has been used as a marker to identify the most suppressive regulatory T cells in humans^23^. In recent years, many glycans have been recognized as active players in the regulation of immune receptor signaling^24–27^. N-aceytllactosamine (Galβ1,4GlcNAc; LacNAc), a universal disaccharide unit of the cell surface, is one of them. Human cytolytic T lymphocyte (CTL) clones exhibit a decrease in their effector activity due to the binding of poly-LacNAc on the N-linked glycans of T-cell receptor (TCR) to extracellular galectin-3, which blocks colocalization of TCR and CD8^24–26^. In addition, there are other reports of LacNAc’s role in modulating immune receptor signaling, but most of them are fragmental^28–33^. One of the reasons behind this issue is that the glycosylation of immune cells is heterogenous and constantly undergoing dynamic changes in a spatial and temporal-dependent manner. Therefore, we argue that adding the information of glycosylation statuses to single-cell multimodal omics would facilitate a better understanding of the relationship between cellular phenotype and cell states, especially for cells in the immune system. However, cell surface glycan is controlled and assembled by multiple processing enzymes. Thus, it is hard to predict the structure and quantity of cell-surface glycans only by the transcription information of relevant processing enzymes by scRNA-seq (**Scheme 1**).

Herein, using a fucosyltransferase-based chemoenzymatic glycan labeling strategy^34, 35^, we designed a new approach for single-cell profiling of the dynamic changes of the relative amounts of LacNAc on cell surface. This method utilizes a recombinant Helicobacter pylori α1,3 fucosyltransferase (FucT), with remarkable donor substrate tolerance, to transfer a C-6 ssDNA-tagged fucose residue to the 3-OH of N-acetylglucosamine (GlcNAc) of the LacNAc disaccharide. Quantitative detection of the LacNAc level is subsequently afforded using established scRNA-seq platforms, simultaneously with transcripts, epitopes or TCR sequence. Using LacNAc-seq, we generated the datasets of single-cell transcriptomes integrated with LacNAc densities in human PBMCs and murine TILs. Further bioinformatic analysis led to the discovery that upregulation of cell-surface LacNAc reflects the upregulation of glycolysis in CD8^+^ T cells. The direct measurement of glycan via chemoenzymatic labeling in scRNA-seq opens up the door of charactering the role of specific glycan via integrated analysis of the connection between specific glycan and thousands of gene transcriptions.

## Results and discussion

### The overview of chemoenzymatic LacNAc labeling-based LacNAc-seq

Integrating the cell-surface LacNAc detection with commercial scRNA-seq platforms requires the labeling of LacNAc with a DNA barcode that converts the amount of LacNAc into a DNA readout for direct measurement using next-generation sequencing (NGS). Previously, we developed a chemoenzymatic method for LacNAc detection, which exploits the donor substrate promiscuity of FucT to transfer a clickable or fluorescent fucose residue for subsequent analysis^34, 35^. Due to the unprecedented promiscuity of FucT, an unnatural GDP-fucose analog bearing a single-stranded DNA can serve as the substrate as well, which, in principle, could be employed for the development of LacNAc-seq. To test this hypothesis, we synthesized GDP-fucose conjugated single-stranded DNAs (GDP-fucose-ssDNA) as the reporter to quantify the level of LacNAc via sequencing. GDP-fucose-ssDNA probes were constructed via the inverse electron-demand Diels-Alder reaction (IEDDA) (**Fig. 1A**), in which the bioorthogonal reaction handle trans-cyclooctene (TCO) with a PEG linker was installed onto the 5’ end of DNA barcodes via the standard amine-coupling procedures. Subsequently, ssDNA barcodes bearing TCO were reacted with GDP-fucose-tetrazine (see synthetic procedures in **Fig. S1**) to afford GDP-fucose-ssDNA (**Fig. 1A**, **Fig. S2**). To confirm GDP-fucose-ssDNA could be transferred to LacNAc on complex glycans, we used free LacNAc and biantennary N-glycan (NA2F) as model substrates to react with GDP-fucose-ssDNA under FucT mediated enzymatic synthesis. Mass analysis of products confirmed the precise fucosylation of LacNAc on different acceptor substrates (**Fig. 1B, 1C**). We also established the conditions to label cell-surface LacNAc with GDP-fucose-ssDNA and identified the optimal concentration in the labeling system (**Fig. S3A**). Flow cytometry analysis revealed that saturation labeling is achieved by using 20 μM of GDP-fucose-ssDNA probes and more than 95% of labeling sites could be blocked by FucT mediated natural fucosylation that used GDP-Fucose as donor substrates (**Fig. S3B**). The stability of ssDNA tagged on LacNAc was also verified, showing no obvious exchange or decay of ssDNA on cell surface within 4 hours (**Fig. S3C, S3D**), which makes it suitable for scRNA-seq. For the adaptation to commercial sequencing platforms, we designed ssDNA barcodes for LacNAc-seq that include a 3’capture sequence, a 36-bp sample barcode and a 5’ PCR handle necessary for library preparation, which is referred as LacNAcTag hereafter. To pursue LacNAc-seq, single cells labeled with LacNAcTag and single RNA capture magnetic beads are seeded into the same well or the same droplet. After cell lysis in wells or droplets, LacNAcTag and mRNA can be simultaneously captured by the capturing magnetic beads functionalized with oligo-dT and are indexed by a shared cellular barcode during reverse transcription. Next, the cDNAs and LacNAcTags are separated and converted into NGS required libraries for subsequent sequencing and data processing (3’LacNAc-seq workflow in **Fig. 1D**).

**Figure 1.**
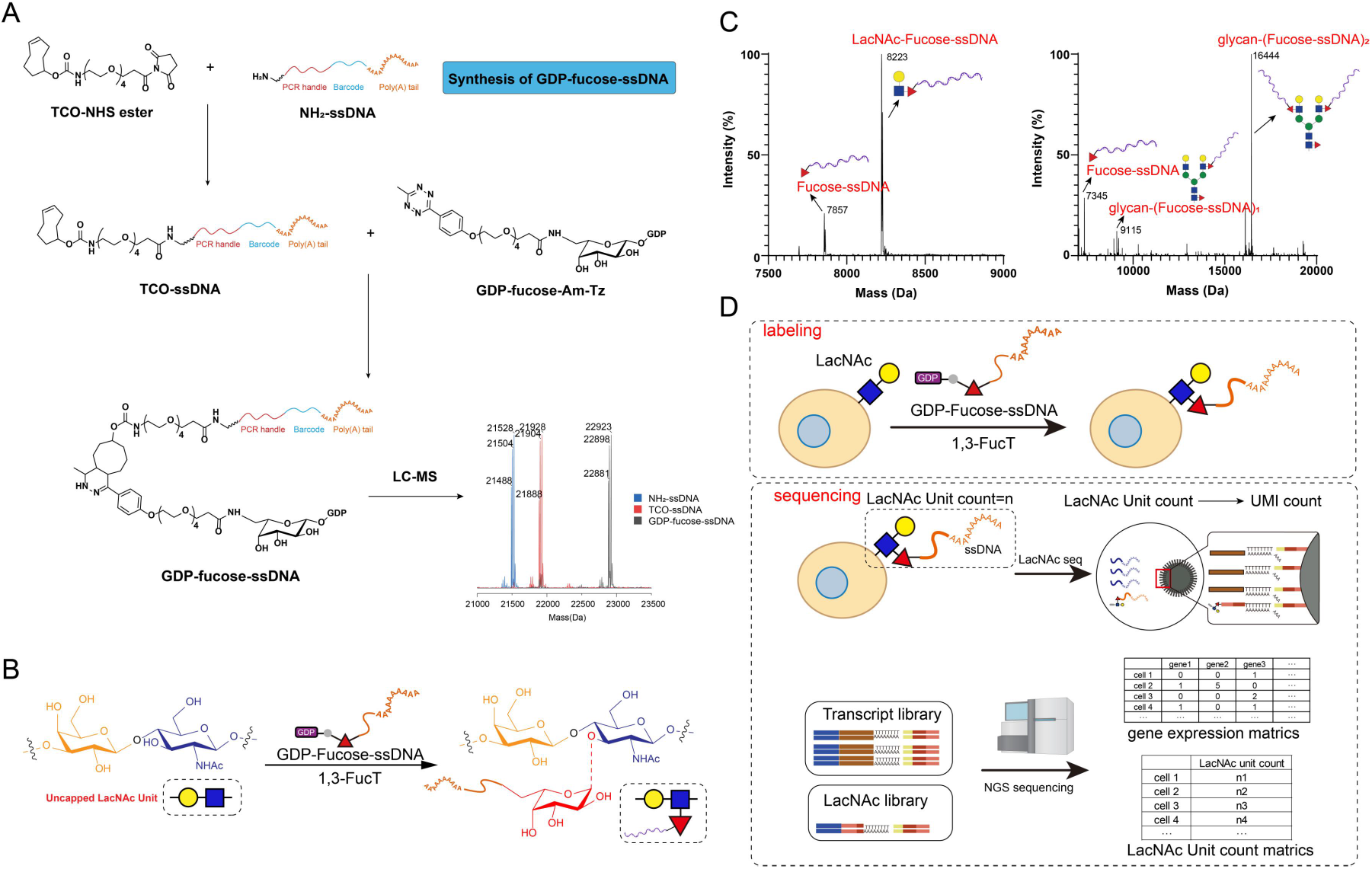
Chemical basis of LacNAc-seq detection. (A) Chemoenzymatic synthesis and LC-MS characterization of GDP-fucose-ssDNA probes. Deconvoluted mass spectra of NH2-ssDNA, TCO-ssDNA and the correlated final product of GDP-fucose-ssDNA are shown. (B) FucT-enabled chemoenzymatic labeling of LacNAc with a GDP-fucose-ssDNA. (C) FucT-enabled transfer of Fuc-ssDNA to LacNAc and biantennary N-glycan (NA2F) was confirmed by mass spectrometry. (D) Schematic overview of the LacNAc-seq workflow in high-throughput scRNA-seq platforms.

### Simultaneous detection of cell-surface LacNAc and transcriptome in single cells

To assess the feasibility of LacNAc-seq on the simultaneous detection of cell-surface LacNAc and transcriptome in single cells, we performed a proof-of-concept experiment using a cell mixture consisting of NK92, Jurkat and A549 cells (**Fig. 2A**). Following a standard scRNA-seq data analysis pipeline, 12343 cells were detected, and the detected cells were clearly clustered into three main groups based on cellular transcriptome. Cell type annotation was then performed accordingly using the transcripts of the known identity markers of NK92, Jurkat and A549 cells (**Fig. 2B**). Using a modified computational workflow for CITE-seq (cellular indexing of transcriptomes and epitopes by sequencing)^14^, we pseudo-aligned LacNAcTag reads to the index of barcodes, which enables the quantification of the LacNAc level of each cell. The results accurately verified the experimental design with a high correlation between unique molecular identifier (UMI) counts for transcripts and LacNAc Tags. Jurkat and A549 cells were found to have comparable LacNAc level, which were much lower than that in NK92 cells (**Fig. 2C**). To validate the quantification results of LacNAc-seq, we analyzed cell-surface LacNAc densities of these three cell lines by flow cytometry, which was enabled by FucT-mediated GDP-fucose-biotin labeling (**Fig. 2A**). Flow cytometric results from biotin tagged LacNAc analysis confirmed the results obtained from LacNAc-seq (**Fig. 2C**). With the correlated quantification of LacNAc barcodes and other transcripts of every single cell on hand, we tried to analyze the LacNAc level with the transcription activity of relevant glycosyltransferases (GTs) that are directly involved in LacNAc synthesis. However, compared to reference proteins, e.g., actin or GAPDH, all GTs of interest have no detectable level of transcriptions, preventing the confirmation of the positive correlation between LacNAc and these GTs (**Fig. 2D**). As a matter of fact, the low transcription of GTs has made it elusive for detection in different scRNA-seq platforms (**Fig. S4**). Nevertheless, the amount of LacNAc is not positively correlated with the transcription level of actin or GAPDH, suggesting that it is not correlated with the size of cell (**Fig. 2D**). By contrast, gene ontology (GO) enrichment analysis indicated that the cell-surface level of LacNAc is upregulated during cell proliferation or division pathways, such as transcription or translations (**Fig. 2E**).

**Figure 2.**
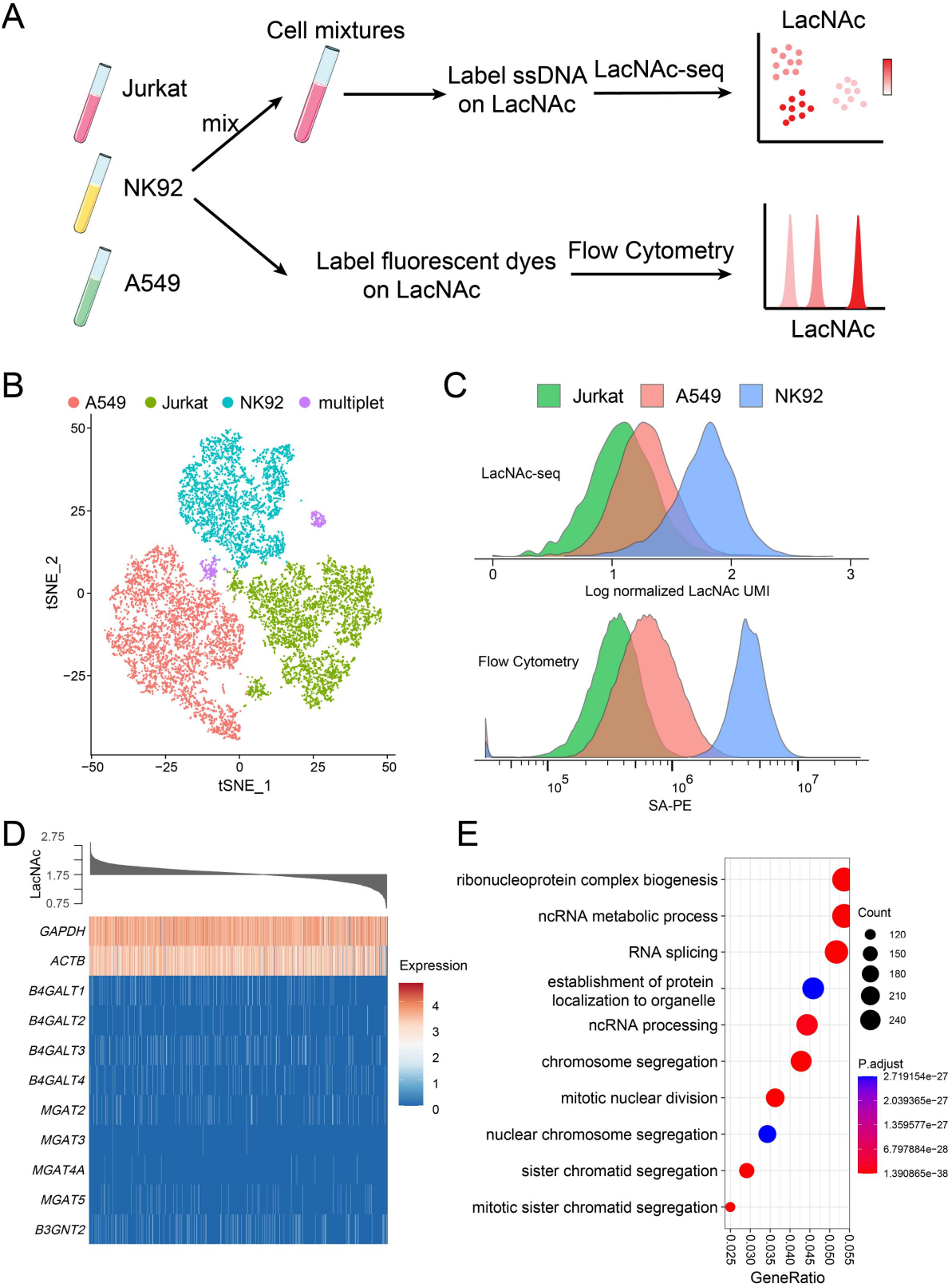
LacNAc-seq enables the simultaneous measurement of transcriptome and cell-surface LacNAc at the single-cell level. (A) Cell mixtures of Jurkat, NK-92, A549 cell lines were barcoded with GDP-fucose-ssDNA probes and then subjected to scRNA-seq. In parallel, each cell line was also labeled with GDP-fucose-biotin probe, followed by streptavidin-APC staining for quantification by flow cytometry. (B) t-SNE plot showing transcriptome-based clustering of 12343 cells from cell mixtures described in A. Distinct cell populations were assigned according to transcriptions of known markers. (C) Merged histograms showing LacNAc density of Jurkat, NK-92, A549 cell lines measured by LacNAc-seq (up) and flow cytometry (bottom). Experimental procedures are shown in A. (D) Heatmap showing the relationship between LacNAc density and expression levels of *GAPDH, ACTB* and GTs related to LacNAc synthesis in NK92 cells. (E) GO analysis revealed biological process enrichment of up-regulated transcripts in the 30% with the highest LacNAc density compared to the 30% with the lowest LacNAc density in NK92 cells.

### Chemoenzymatic LacNAc labeling has minimal influence on the transcriptional status of immune cells

It has been reported that methods for probing cell-surface glycans may affect the physiological status of cells due to the cross-linking of receptors through glycosylation^36^. To assess the perturbation of chemoenzymatic labeling on immune cells, we performed scRNA-seq to analyze the fresh mouse thymocytes labeled with FucT and GDP-fucose-ssDNA (**Fig. 3**). In parallel, untreated thymocytes and thymocytes labeled by ssDNA-conjugated Phytohemagglutinin-L (L-PHA), a plant lectin that binds to the similar structure of LacNAc, were used as controls. Through analyzing the scRNA-seq data of these three samples, we found that the chemoenzymatic labeled group displayed the similar transcriptome-dependent unsupervised clustering of immune cells as the untreated group (**Fig. 3D, 3E**), while the L-PHA-stained group showed an increased population of proliferating T cells (**Fig. 3D, 3E**), suggesting that the L-PHA-based labeling induce T-cell activation via known mechanisms of binding and crosslinking of T cell receptors^36^. Moreover, principal-component analysis (PCA) of single-cell-based transcriptomes of these three groups clearly showed that the chemoenzymatic labeling group clustered together with the untreated group and both these two groups clustered separately from the L-PHA labeled group (**Fig. 3F**). Further analysis of the differentially expressed transcripts revealed that the upregulated transcripts in L-PHA-stained group were mainly correlated with cell growth and division (**Fig. 3G, Fig. S5**). According to these results, we speculate that the tetrameric L-PHA likely induces the crosslinking of certain cell-surface receptors and triggers the downstream signaling, even at low temperature. By contrast, the chemoenzymatic labeling does not induce any cross-linking to alter cell’s properties, and therefore it serves as a better approach than the lectin labeling in transcriptome-based single-cell multiomics.

**Figure 3.**
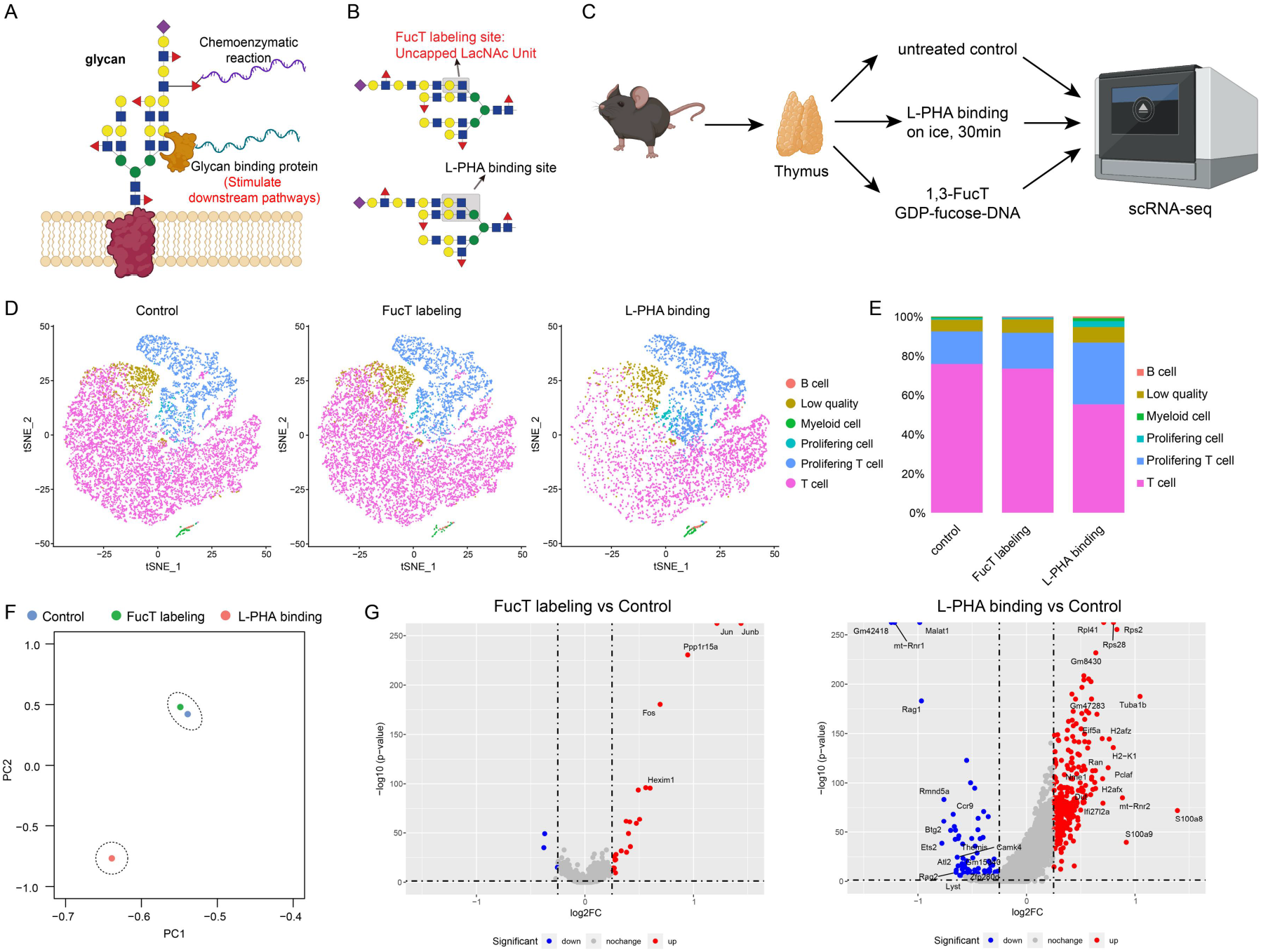
Chemoenzymatic labeling of cell-surface LacNAc with ssDNA does not change the ratio and status of immune cells. (A) Glycan-binding protein and chemoenzymatic glycan editing-enabled methods for introducing ssDNA probes onto LacNAc residues for single-cell analysis. (B) Unique sites of N-glycans recognized by L-PHA and FucT. (C) Fresh mouse thymocytes were labeled with FucT/GDP-Fuc-ssDNA or L-PHA-ssDNA and then subjected to scRNA-seq to analyze the transcriptional changes when compared with the untreated sample. (D) tSNE plot showing transcriptome-based clustering of single-cell expression profiles of the three samples. (E) Cell compositions in mouse thymocyte samples following different labeling procedures. (F) PCA analysis revealed transcriptional differences among control, FucT labeling and L-PHA staining. (G) Volcano plot showing differential transcripts between FucT labeling, L-PHA binding and the untreated groups.

### Single-cell human PBMC LacNAc map revealed by LacNAc-seq

To determine if LacNAc-seq is applicable to human samples and can be combined with CITE-seq, we performed LacNAc-seq combined with CITE-seq using a mixed PBMC sample that contained freshly isolated PBMCs and PBMCs that were activated by anti-CD3/CD28 overnight (**Fig. 4A**). CITE-seq relies on DNA-barcoded antibodies to convert the detection of proteins into a quantitative, sequenceable readout (**Fig. 4B**). As expected, independent unsupervised analysis of the transcriptome and epitopes data revealed consistent cell classifications except for the CD4 epitope, which was partially blended when analyzing the transcriptome but clearly identified in CITE-seq epitope channel by the DNA barcoded CD4 antibody (**Fig.4B, Fig.S6A**). In parallel, LacNAcTag was quantified together with that of the CITE-seq epitopes and clearly assigned to each cell population (**Fig.4B**), demonstrating the compatibility of the LacNAc-seq procedure with CITE-seq. Mapping LacNAcTag on the t-SNE plot revealed that immune cells in PBMCs display heterogeneous LacNAc densities, in which monocytes have higher LacNAc densities than T and B cells (**Fig. 4C, 4D, Fig. S6B**). Previous studies have provided evidence that T-cell activation is accompanied by dramatic changes of cell-surface receptor glycosylation patterns. For example, upon T-cell activation, an increase of β1,6-GlcNAc branched N-glycosylation of CTLA-4 takes place, which augments CTLA-4 retention at the T cell surface, thereby suppressing T cell activation promoting immune tolerance^37, 38^. To gain a better understanding if CD8^+^ T cells at different activation status may express different levels of LacNAc, we filtered *CD8A* positive cells in the mixed PBMC sample and annotated them as CD8^+^ T cells, from which five clusters were identified via unsupervised clustering analysis (**Fig. 4E, Fig. S7**). These clusters were further annotated as naïve, early activated, effector, effector memory and central memory CD8^+^ T cells, respectively. We found that effector and effector memory CD8^+^ T cells displayed comparable LacNAc densities, which were higher than that of naïve CD8^+^ T cells. Interestingly, central memory CD8^+^ T cells were found to have similar levels of LacNAc as naïve CD8^+^ T cells presumably due to their similar quiescent statuses (**Fig. 4F**). We compared the transcript differences between Lac.hi (LacNAc level in top 30%) and Lac.low (LacNAc level in bottom 30%) in CD8^+^ T cells, and found that the transcripts positively associated to Lac.hi were enriched in many transcripts related to T cell activation and effector function, such as *IFNG, GZMB,* etc (**Fig. 4G**). To infer the dynamic changes of LacNAc level during CD8^+^ T cell differentiation, we used Monocle algorithm to perform pseudotime analysis on the CD8^+^T cell subsets (**Fig. 4H**). These analyses revealed a trajectory originating at naive and central memory CD8^+^ T cells, through the early activated CD8^+^ T cells, and gradually evolves into effect and effector memory CD8^+^ T cells. The level of LacNAc in pseudotime gradually increased along the differentiation trajectory (**Fig. 4H**), indicating that the cell-surface LacNAc is upregulated during the T cell activation process. Taken together, these results suggest that the level of cell-surface LacNAc may serve as a general CD8^+^ T cell activation marker.

**Figure 4.**
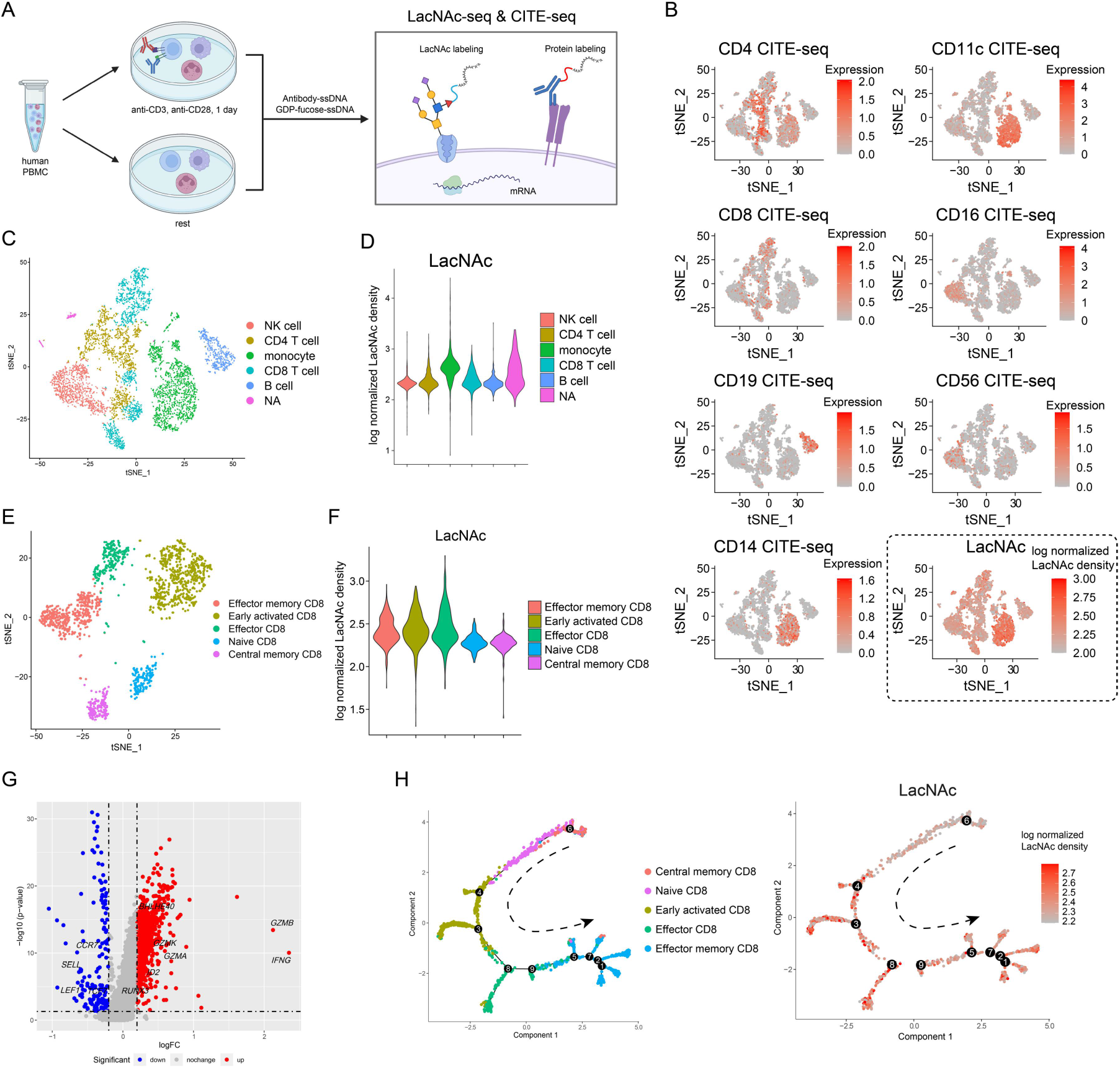
Simultaneous detection of the transcriptome, cell surface protein expression and LacNAc in the mixtures of activated and rested human PBMCs. (A) Schematic illustration of the workflow for detection of LacNAc, transcriptome and cell surface protein expression. PBMCs from healthy donors were divided into two samples. One was stimulated with anti-CD3 and anti-CD28 antibodies for one day, while the other was kept rested for one day. After that, both stimulated and unstimulated PBMCs were mixed as one sample (named mixed PBMCs). (B) tSNE plots showing the expression level of cell surface proteins by CITE-seq and LacNAc density. (C) tSNE clustering of mixed PBMCs by CITE-seq. n = 5865 cells. (D) Violin plot showing LacNAc density on each immune cell population. (E) tSNE analysis and graph visualization of CD8^+^ T cells from mixed PBMC samples. (F) Violin plot showing LacNAc density on each population of CD8^+^ T cells. (G) Volcano plot showing the transcript differences of 30% CD8^+^ T cells with high LacNAc expression vs. 30% CD8^+^ T cells with low LacNAc expression. (H) Left: Monocle-generated plots showing pseudotime ordering and differentiation trajectory of CD8^+^ T cells in mixed PBMCs. Right: Monocle-generated plots showing the change in LacNAc following the trajectory timeline.

### LacNAc on CD8^+^ T cell is associated with T cell differentiation state but not with TCR clonal expansion

As LacNAc is highly upregulated upon T cell activation, we explored whether the dynamic changes of LacNAc are mainly controlled by the activation via the TCR signaling pathway. If this is true, the increased level of LacNAc may indicate the antigen-driven clonal expansion of T cells. This would be quite useful for the rapid identification of tumor antigen-specific T cells from tumor infiltrating lymphocytes (TILs) since antigen-specific T cells are continuously stimulated by tumor antigens to trigger TCR signaling. To test this hypothesis, TILs from a E0771 murine triple negative breast cancer model were isolated. Enriched CD45^+^ cells were labeled with LacNAcTag (see 5’ workflow sequence in **SI Methods**) and then subjected to LacNAc-seq combined with scTCR-seq that allows the simultaneous detection of transcriptomes, T cell clonotypes and LacNAc level (**Fig. 5A**, see the integrated workflow of LacNAc-seq and 10x Genomics 5’ scRNA-seq with scTCR-seq in **Fig. S8A**). Following a standard scRNA-seq data analysis pipeline, 6129 cells were detected and clustered by transcriptomes in an unsupervised manner (**Fig. 5B**). In parallel, the LacNAc level was quantified by the UMI count and normalized logarithmically (**Fig. 5C, Fig. S8B**). Consistent to what was observed in human PBMCs, varying LacNAc densities were found in different immune cell types, in which monocytes and macrophages have highest LacNAc level (**Fig. 5C**). Within a specific cell type, variations in the LacNAc level were also observed. CD8^+^ T cells (*CD8A* positive) extracted from total TILs were then analyzed by the dimension-reduction and unsupervised clustering methods (**Fig. 5D, Fig. S9**). As shown in the violin plot of LacNAc quantification, the violin plot of LacNAc quantification, effector CD8^+^ T cells have significantly higher LacNAc levels than both exhausted and naïve or memory CD8^+^ T cells, while exhausted CD8^+^ T cells exhibited a slightly higher LacNAc level than naïve or memory CD8^+^ T cells (**Fig. 5E**). To study the relationship of T cell clonal expansion and the LacNAc level on CD8^+^ T cells, we projected TCR clonotypes and their expansion fold into the tSNE plot of CD8^+^ T cell clustering (**Fig. 5F**). However, there was no significant correlation between the LacNAc level and T cell clonal expansion (**Fig. 5G**). As revealed by the pseudotime analysis of four CD8^+^ T cell subsets via Monocle algorithm, the LacNAc dynamic change has no obvious overlap with TCR clonal expansion (**Fig. 5H, 5I, 5J**). Previous studies by Bevan and coworkers revealed that there is a positive correlation between T-cell clonal expansion and the affinity of the priming ligand^39^. To confirm that LacNAc level is not governed by T cell priming strength, we used a Listeria-Ova (Lm-Ova) infection model to determine the LacNAc densities of T cells that are stimulated under strong and weak TCR engagers, respectively^39^. We adoptively transferred OT-I T cells into two sets of recipient mice and infected mice with Lm-Ova-N4 (SIINFEKL, the WT sequence, strong) and Lm-Ova-T4 (SIITFEKL, single amino acid substituted altered peptide ligand, weak), respectively. As expected, we observed lower expansion of OT-I cells in response to Lm-Ova-T4 vs. Lm-Ova-N4 infection (**Fig. 5K**). By contrast, LacNAc densities were found to be identical on OT-I CD8^+^ T cells in both groups, which is 8 times more than that of the host naïve T cells (CD44^lo^CD62L^hi^) (**Fig. 5K**). Taken together, these results confirm that the LacNAc level is upregulated upon T cell activation and is not correlated with the TCR signaling strength.

**Figure 5.**
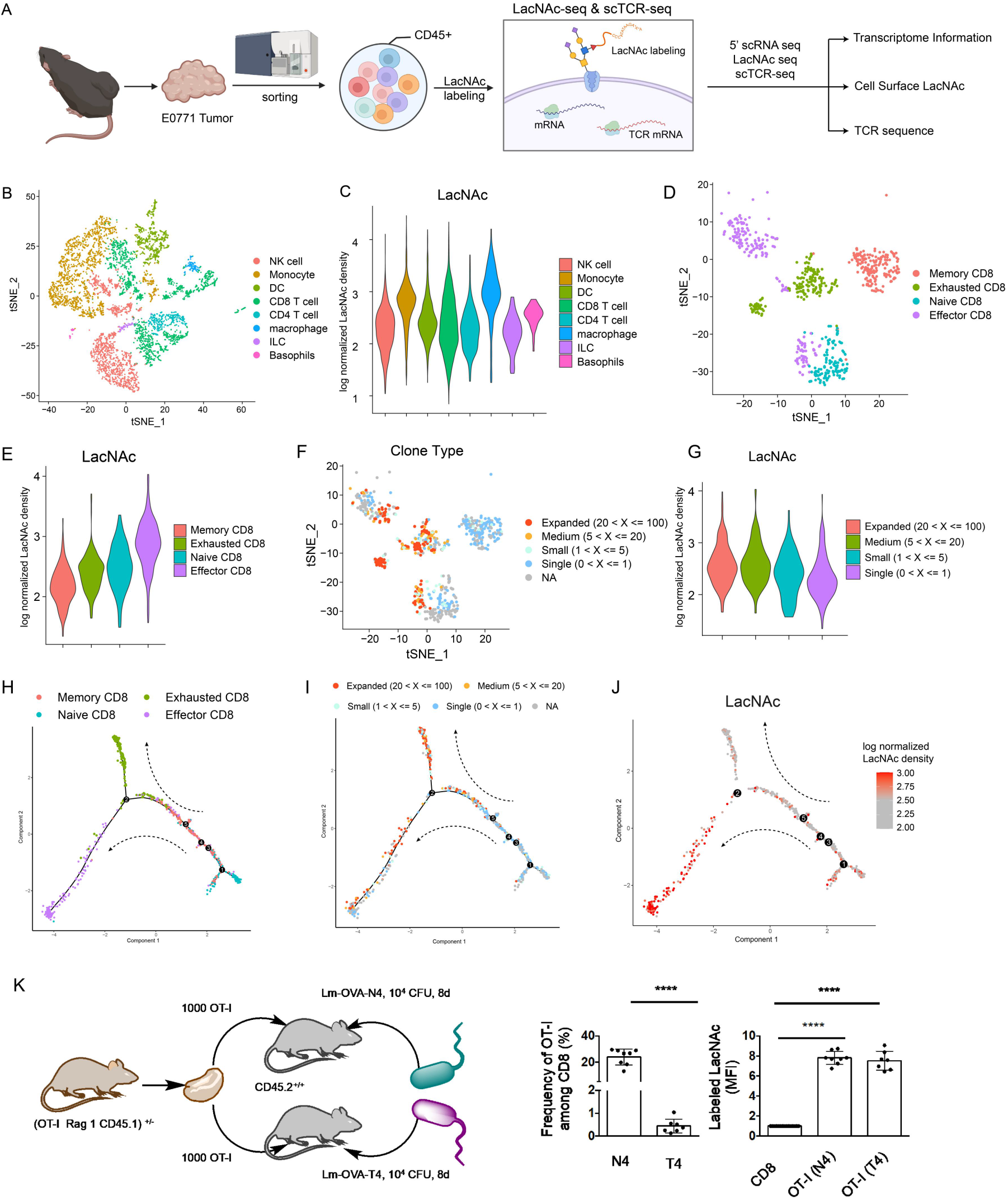
LacNAc density on CD8^+^ T cell is related to the T cell differentiation state. (A) The workflow of 5’ LacNAc-seq and scTCR-seq on E0771 tumor infiltrating lymphocytes (TILs). E0771 TILs (CD45^+^) were isolated from E0771 tumor on day 14. To perform 5’LacNAc seq and scTCR seq on the 10x genomics platform, we followed the rule of Biolgend TotalseqC™ type oligo sequence in making GDP-fucose-ssDNA probes. Transcriptomes, TCR sequence and LacNAc density were detected simultaneously in single cells. (B) tSNE analysis and graph visualization of E0771 TILs. n=6129 cells. (C) Violin plot showing the LacNAc density on each population of TILs. (D) tSNE analysis and graph visualization of CD8^+^ T cells in E0771 TILs. (E) Violin plot showing the LacNAc density on each population of T cells. (F) Expanded CD8^+^ T cells clones projecting across the tSNE clustering map shown in D. Each dot denotes an individual T cell; colour denotes the fold of T cell clonal expansion. (G) Violin plot showing the LacNAc expression across CD8^+^ T cell clusters divided by TCR clonal expansion shown in F. (H) Monocle-generated plots showing pseudotime ordering and differentiation trajectory of CD8^+^ T cells in E0771 TILs. (I) Expanded CD8^+^ T cells clones projecting across the Monocle-generated plots. (J) Monocle-generated plots showing the change of LacNAc expression on CD8^+^ T cells following the trajectory timeline. (K) Splenocytes from 8d-infected mice are divided into endogenous and OT-I, N4- or T4-responsive T cells. Flow cytometric analysis of LacNAc expression and cell frequency are shown.

### CD8^+^ T cell-surface LacNAc level is positively correlated with the glycolytic activity

The observation that the activated, effector CD8^+^ T cells had higher LacNAc level than naïve T cells, combined with the finding that exhausted and naïve or memory CD8^+^ T cells had similar LacNAc level, prompted us to further explore the mechanism that regulate the LacNAc level on CD8^+^ T cells. As glycan biosynthesis is known to be regulated by cellular metabolism^40^, we used gene set variation analysis (GSVA) to calculate the pathway activated score of glycolysis^41^, TCA cycle and oxidative phosphorylation, and compared the difference between Lac.hi (LacNAc level in top 30%) and Lac.low (LacNAc level in bottom 30%) in CD8^+^ T cells of E0771 TILs (**Fig. 6A**). As revealed in the heatmap, the population of Lac.hi displayed the obvious upregulated metabolic activity, especially in glycolysis, when compared to the Lac.low population. We further performed a cellular metabolism-focused GSEA analysis of Lac.hi vs. Lac.low cells. The results indicated that the upregulation of glycolysis is much more significant than the other two pathways (**Fig. 6B**). Moreover, the dynamic glycolysis activity profile during CD8^+^ T-cell activation is positively correlated with the dynamic changes of cell-surface LacNAc level (**Fig.6C**). We also performed the same analysis of CD8^+^ T cells from the mixed human PBMC sample that contain both quiescent and activated T cells (**Fig. S10**). Consistent to what we had observed for TILs, the glycolysis process was always enriched in the high LacNAc level group when CD8^+^ T cells were analyzed in total or in subgroups, whereas no such relationships were found for oxidative phosphorylation and TCA cycle with LacNAc densities (**Fig. S10C, Fig. S10D**). These observations implicate that glycolysis may play a critical role in regulating LacNAc biosynthesis. Several intermediate products in glycolysis serve as raw materials for the assembly of cell-surface glycans. For example, fructose-6-phosphate (F-6-P) enters the hexosamine pathway to produce UDP-GlcNAc for the biosynthesis of LacNAc^27^ (**Fig. 6D**). To further elucidate the relationship between the LacNAc level and the glycolytic activity in specific CD8^+^ T cell subsets, we analyzed public databases that contains bulk RNA seq data of transcript expressions related to glycolysis and LacNAc biosynthesis of CD8^+^ T cells in different differentiation states^42^ (**Fig. 6E**). As well documented, naïve CD8^+^ T cells are metabolically quiescent. They uptake a basal level of nutrient and employee OXPHOS as their primary pathway of ATP production. Upon antigen encounter, CD8^+^ T cells are activated, which is accompanied by an increase of nutrient uptake and glycolytic and glutaminolytic metabolism. Transition to the memory stage is characterized by a return to quiescent metabolism with reduced glycolysis and increased reliance on FAO to fuel OXPHOS^43^. Consistently, compared with naïve and memory CD8^+^ T cells, effector CD8^+^ T cells generally expressed higher levels of transcripts involved in glycolysis and LacNAc biosynthesis according to the analysis of public data (**Fig. 6E**). First, *Hk1* (encoding Hexokinase 1, HK) and *Gpi1* (encoding glucose-6-phosphate isomerase 1, GPI1) were upregulated in effector CD8^+^ T cells, which are responsible for catalyzing the conversion of glucose to fructose-6-phosphate (F-6-P), supplying materials for both of glycolysis and hexosamine synthesis pathways (**Fig. 6D**). Furthermore, the most upregulated transcripts in effector CD8^+^ T cells also included *Pfkp* (encoding Phosphofructokinase, PFK), *Ldha* (encoding Lactate Dehydrogenase A, LDH), *Gfpt1* (encoding Glutamine-Fructose-6-Phosphate Transaminase 1, GFPT), *B4galt1* (encoding Beta-1,4-galactosyltransferase 1, B4GALT1) and *Mgat2* (encoding Alpha-1,6-Mannosyl-Glycoprotein 2-Beta-N-Acetylglucosaminyltransferase, MGAT2)^44^ (**Fig. 6D**). Among them, PFK and LDH are key players in glycolysis that catalyze the further glycolysis of F-6-P to lactate; GFPT is the rate-limiting enzyme that catalyzes the conversion of F-6-P to glucosamine-6-phosphate to enter the hexosamine pathway; B4GALT1 and MGAT2 are key glycosyltransferases that assembly LacNAc (**Fig. 6F**). Likewise, compared to effector CD8^+^ T cells, exhausted CD8^+^ T cells exhibit repressed glycolytic and mitochondrial metabolism^45^. As shown in **Fig. 6E**, although exhausted CD8^+^ T expressed high levels of HK which is required for producing high levels of glucose-6-phosphate (G-6-P), all of the other transcripts related to glycolysis and N-glycan biosynthesis were expressed at a relative lower level as compared to effector CD8^+^ T cells, e.g., GPI1, B4GALT1, MGAT1-4. Indeed, exhausted CD8^+^ T cells were found to express lower LacNAc level than effector CD8^+^ T cells (**Fig.5E, Fig. S11**). Together, these findings reveal that LacNAc level is most upregulated in effector CD8^+^ T cells that possess the highest glycolytic activity among all stages of CD8^+^ T cells. Therefore, high LacNAc densities, in turn, may indicate the metabolic state of CD8^+^ T cells that maintain a high level of glycolysis.

**Figure 6.**
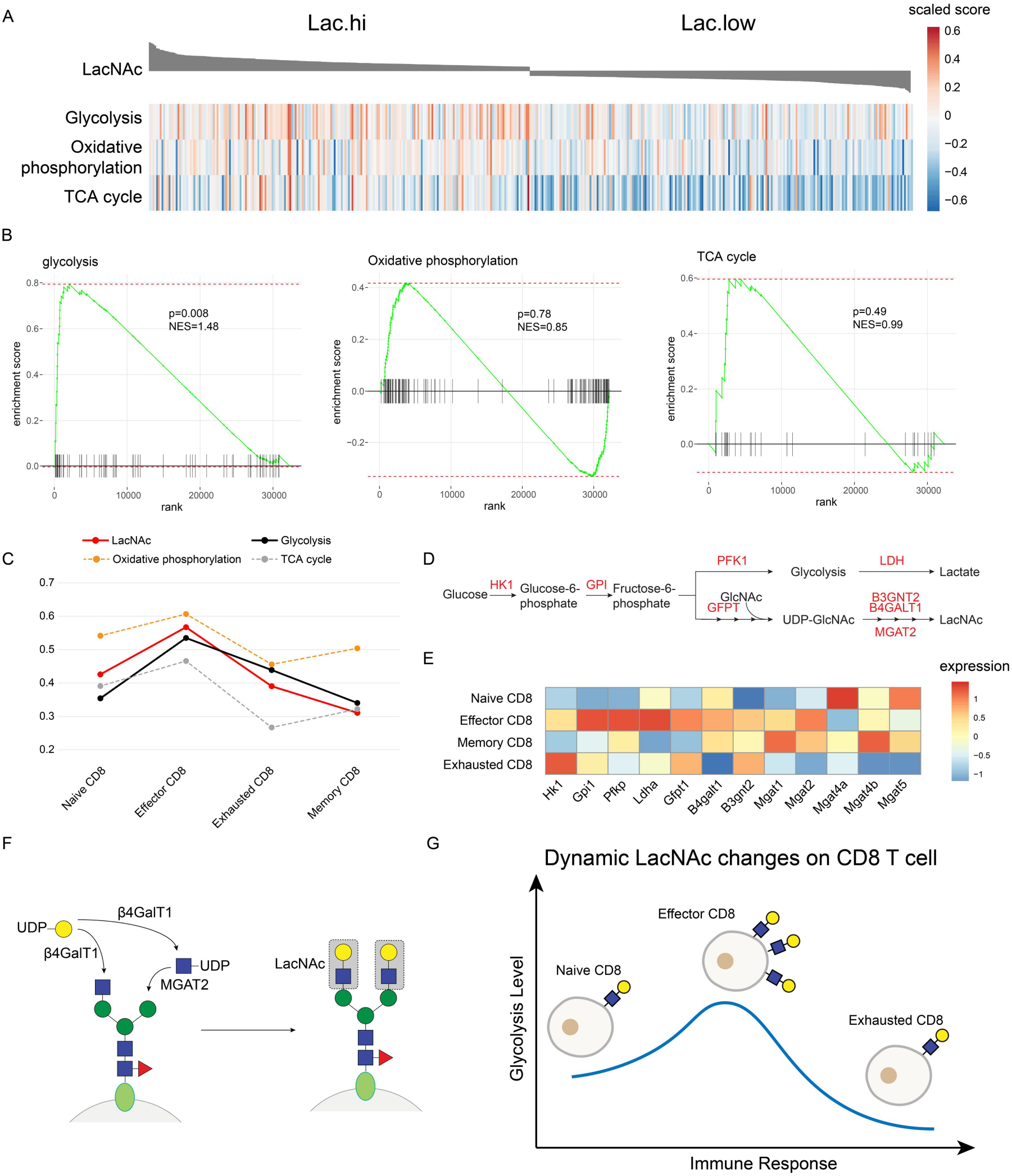
LacNAc density on CD8^+^ T cell reflects the metabolic state of upregulation of glycolytic activity. (A) Heatmap showing the CD8^+^ T cell Gene Set Variation Analysis (GSVA) based pathway activity scores for glucose metabolism pathways, visualized for the top 30% and bottom 30% of LacNAc density. The pathway activity scores were scaled by row. (B) Gene set enrichment analysis (GSEA) of glycolysis, TCA cycle and oxidative phosphorylation for the comparison of transcripts of Lac.hi versus Lac.low. (C) Line plot showing cluster-averaged GSVA-based glucose metabolism pathways activity score and LacNAc density following CD8^+^ T cell activation. Data was scaled to 0-1. (D) Overview of glycolysis and its branching pathway of N-glycosylation. Fructose-6-phosphate produced by glucose is the key starting material for both of glycolysis and LacNAc synthesis. Key enzymes are marked in red. (E) RNA-seq heatmap presenting differentially expressed transcripts of glycolysis and N-glycan biosynthesis (raw data adapted from GSE88987) among naive, effector, memory and exhausted CD8^+^T cells. (F) Schematic illustration showing key enzymes of LacNAc biosynthesis. (G) Dynamic LacNAc changes on CD8^+^ T cell surface as an indicator of their glycolysis states, of which T cells under activated states upregulate LacNAc level, as compared to naive or exhausted states.

### Summary and outlook

Multimodal single-cell omics have provided powerful tools to discover, characterize, and catalog cell types and states especially for studying the immune system beyond what can be achieved by employing transcriptomics alone, which usually cannot separate clusters of immune cells with different functions due to similar gene-expression profiles^11^. Although glycosylated molecules play essential roles in immune cell differentiation and trafficking, due to the low transcription levels of glycosyltransferases and the lack of tools for glycan decoding, scRNA-seq rarely provides insightful glycosylation-related information.

In this study, we developed LacNAc-seq based on chemoenzymatic glycan editing that was combined with CITE-seq and scTCR-seq to simultaneously detect cell-surface LacNAc and phenotypic markers, transcripts, and T-cell clonotypes. Featuring scalability, wide adaptability in multiple platforms, and specificity in glycan recognition, LacNAc-seq realized the first single-cell LacNAc mapping of human immune cells. We discovered that the LacNAc level is correlated to the activation state of immune cells, especially that of CD8^+^ T cells. Interestingly, via GSEA analysis we found that high LacNAc level reflects an activated state with high glycolytic metabolism in CD8^+^ T cells. This observation is further confirmed by the analysis of the RNA-seq data of murine T cells (from the Gene Expression Dataset GSE88987). CD8^+^ T-cell up-regulate glycolysis upon the stimulation by antigens or other activation signals, providing F-6-P as the raw material for the synthesis of LacNAc. In parallel, antigen-stimulation promoting the expression of LacNAc biosynthesis-related enzymes, such as B4GALT1, B3GNT2 and MGAT2, which lead to an increase of LacNAc level on the cell surface. As a result, high LacNAc levels might serve a unique indicator to identify the high glycolytic activity and effector function of CD8^+^ T cells accordingly (**Fig. 6G**). Interestingly, these LacNAc-bearing glycoproteins could be recognized and targeted by cancer cells through the secretion of LacNAc binders, etc. Galectin-3, which contribute to the TIL dysfunction in the tumor microenvironment through breaking the physical dissociation between TCR and CD8, or even inducing T cell apoptosis^24–26^. Our new findings, combined with these previous observations, indicate that LacNAc as a cell-surface glycolysis marker could only be applied to specific T cell subtypes (i.e., CD8^+^).

Similar to CITE-seq, chemoenzymatic glycan labeling-enabled glycan-seq is inherently customizable and scalable. Moreover, the direct transfer of ssDNA barcode to glycosylated receptors will not induce crosslinking of cell surface receptors and subsequent signaling transduction. By contrast, other detection methods using lectins or avidins that have multivalent binding sites potentially have the issue of changing cell status via crosslinking receptors. Although chemoenzymatic glycan labeling with DNA barcode are very compatible with glycan-seq, glycosyltransferases that have flexible donor substrate scopes to incorporate probe molecules, e.g., FucT, are quite limited^46,47^. Directed evolution may provide a solution for providing more glycosyltransferases with high efficiency, flexible donor scope, and precise acceptor recognition to expand the toolkit of chemoenzymatic glycan-seq for single cell applications. We envisage that glycan-seq would be easily integrated with other recently developed high-throughput multimodal single-cell sequencing techniques, e.g., ATAC with Select Antigen Profiling by sequencing (ASAP-seq)^18^, single-nucleus chromatin accessibility and mRNA expression sequencing (SNARE-seq)^48^, Perturb-seq^49^. And these techniques could further be empowered by novel computational methods to provide rich resource at the single-cell resolution for the study of cell identity and function in the future.

## Author Contributions

J.P.L, P.W., W.Y. and A.S.J designed experimental strategies. W.Y., X.Z., A.S.J., Y.Y. and Z.Z. performed experiments. W.Y., X.Z., J.P.L., and P.W. prepared the figures and performed bioinformatics analysis. J.P.L., P.W., and W.Y. wrote the manuscript. All authors analyzed the data and edited the manuscript.

## Notes

All scRNA-seq experiments were performed in Nanjing University, China.

## ACKNOWLEDGMENT

J.P.L. acknowledges the support from the National Key R&D Program of China (2019YFA09006600), National Natural Science Foundation of China (21977048 and 92053111), Fundamental Research Funds for the Central Universities (0205/14380225), Beijing National Laboratory for Molecular Sciences (BNLMS202008), Jiangsu Specially-Appointed Professor Plan, Program for Innovative Talents and Entrepreneur in Jiangsu, and the Excellent Research Program of Nanjing University (ZYJH004). P.W. is supported by NIH (R35 GM139643). Z.J.G. acknowledges the support from National Natural Science Foundation of China (21731004 and 91953201) and Natural Science Foundation of Jiangsu Province (BK20202004). We thank Dr. H. Shen generously provide us the Ovalbumin-expressing Listeria monocytogenes (Lm-Ova).

---

**Scheme 1.**
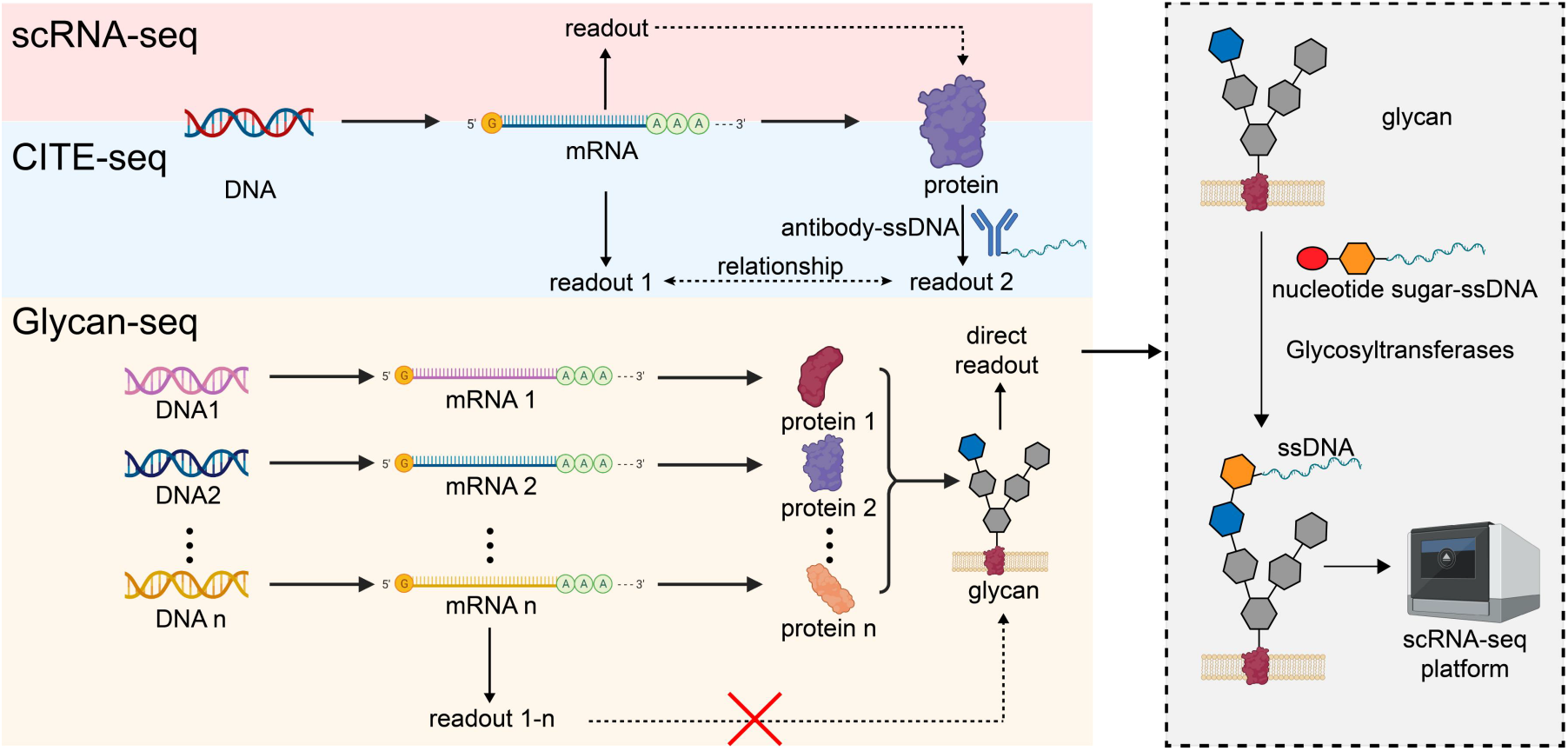
Glycan-seq in single-cell multi-omics. Schematic illustration of the requirement of the direct glycan detection in transcriptome-based single-cell multi-omics. scRNA-seq assumes the transcription level could represent the expression level of protein of interest and uses the transcriptome to define and describe cell status. To correct the difference between transcription and expression, CITE-seq was developed to detect the expression of cell surface epitopes via ssDNA barcoded antibodies, simultaneously with the transcriptome. However, cell-surface glycan status is controlled by multiple enzymes, which could not be predicted only by the analysis of the transcripts of relevant enzymes. Thus, compared to the cell-surface protein epitopes, direct detection of glycan with ssDNA barcoded probes by chemoenzymatic method is much more required in transcriptome-based single-cell multi-omics.

